# CD73-derived adenosine at the blood-brain barrier confers protection in a mouse model of ischemic stroke

**DOI:** 10.64898/2026.06.19.732935

**Authors:** Maria Stamataki, Enrico Maria Costanzo, Julia Lüschow, Johanna Hiefner, Alexander Veltkamp, Kristoffer Riecken, Tobias Mummert, Michael Kaul, Ceren Saygi, Malik Alawi, Anna Worthmann, Björn Rissiek, Tim Magnus, Jakob Körbelin

## Abstract

Ischemic stroke remains a leading cause of death and disability, and current reperfusion therapies do not address the secondary neuroinflammatory response following blood–brain barrier (BBB) disruption. Purinergic signaling critically regulates this process: extracellular ATP promotes inflammation, whereas its enzymatic conversion into adenosine exerts tissue-protective effects. Notably, the ectonucleotidase CD73 (NT5E), which catalyzes AMP-to-adenosine conversion, is highly expressed by human but not murine brain endothelial cells (BECs). Here, we investigated the role of endothelial CD73 in ischemic stroke using an AAV vector engineered for selective transduction of murine BECs to induce BBB-specific CD73 expression. Endothelial CD73 enhanced extracellular ATP degradation toward adenosine generation and established a purine metabolism profile resembling that of human BECs. In the transient middle cerebral artery occlusion (tMCAO) mouse model, BBB-targeted CD73 expression reduced infarct volume by 40% and prevented early mortality within 48 h after reperfusion. Transcriptomic and flow cytometric analyses revealed altered leukocyte responses, including increased recruitment of monocytes/macrophages whose gene expression signatures were consistent with inflammation-resolving programs. These findings identify endothelial CD73 as an important regulator of post-ischemic neuroinflammation and highlight species-specific differences in BBB purine metabolism with implications for translational stroke research.

## INTRODUCTION

Stroke remains a major global health burden, representing one of the leading causes of death and long-term disability worldwide[1]. While reperfusion therapies such as intravenous thrombolysis and mechanical thrombectomy have significantly improved acute management, they apply to a minority of patients and fail to address the secondary neuroinflammatory cascade that contributes to infarct progression[2, 3]. A key component of this inflammatory response is the disruption of the blood-brain barrier (BBB), which permits infiltration of peripheral immune cells and exacerbates tissue injury[4–6]. The BBB is a highly specialized interface formed by brain endothelial cells, pericytes, and astrocytic endfeet. Brain endothelial cells (BECs) play a pivotal role in maintaining CNS homeostasis by regulating molecular and cellular traffic between the bloodstream and the brain parenchyma[7]. Ischemic stroke disrupts BBB integrity, leading to vascular leakage, immune cell infiltration, and further neuronal damage[8]. Purinergic signaling is fundamental in regulating the stroke-induced inflammatory response at the BBB, and all involved cell types express substantial levels of purinergic receptors in different combinations[9–11]. We have shown that transient middle cerebral artery occlusion (tMCAO) leads to the release of high levels of extracellular ATP, which negatively impacts the condition’s outcome[12]. While extracellular ATP promotes inflammatory responses via P2X signaling, it can be degraded by ectoenzymes into adenosine, which exerts anti-inflammatory effects via adenosine receptors[9, 13, 14]. Canonically, the balance of extracellular ATP and adenosine is regulated by the ecto-nucleoside triphosphate diphosphohydrolase 1 (NTPDase1/ CD39) and the ecto-5’-nucleotidase (NT5E/CD73). While CD39 hydrolizes adenosine triphosphate (ATP) to adenosine diphosphate (ADP) and further to adenosine monophosphate (AMP), CD73 degrades AMP to adenosine (ADO). Both enzymes are expressed by a variety of cells and play a fundamental role in shaping the immune response to ischemic injury. Different murine stroke models so far have shown conflicting effects of a global CD73 knockout. While CD73-deficient mice presented with larger infarcts and increased immune cell infiltration in the photothrombotic stroke model[15], we did not detect significant effects of a global CD73 knockout in the tMCAO model[16]. Assuming that CD73 expression would exert different effects on different cell types, we specifically investigated the role of BBB-expressed CD73 in the context of stroke. Surprisingly, the expression of CD39 and CD73 at the BBB differs vastly between humans and mice. While human BECs express substantial levels of CD73, they only express low levels of CD39. Vice versa, CD39 is strongly expressed by murine BECs, while CD73 is almost absent[9]. To specifically assess the importance and therapeutic potential of BBB-expressed CD73 in stroke, in this study, we employed AAV-BR1, an adeno-associated virus capsid engineered to transduce murine brain microvascular endothelial cells after systemic administration[17]. We generated an AAV-BR1 vector encoding murine CD73 under the control of the ubiquitous CAG promoter and used this vector to overexpress CD73 in murine BEC. Indeed, CD73-expressing murine BECs rapidly converted extracellular ATP to adenosine, comparable to human BECs. When assessing the effect of CD73 overexpression in the tMCAO stroke model, vector-treated mice showed a reduced infarct volume by 40% compared to control animals, as assessed by T2-weighted MRI. The reduced infarct volume, in turn, was reflected by an elimination of early mortality within the acute phase, 48 hours after reperfusion (2d survival: <70% vs. 100%). Bulk RNA sequencing of CNS immune cells revealed substantial changes in gene expression. We observed differential upregulation of numerous chemotaxis-related genes and a multitude of genes primarily associated with myeloid cells, several of which were consistent with an inflammation-resolving phenotype. The change in gene expression was accompanied by increased stroke-induced infiltration of monocytes/ macrophages into the brains of mice with CD73-expressing BECs. The apparently protective effect of the increased leukocyte infiltration presumably was brought about by upregulating genes involved in tissue repair and inflammatory balance when comparing AAV-CD73-treated animals to animals treated with an empty control vector, thereby potentially limiting the stroke-induced tissue damage. Taken together, our data show a strongly beneficial effect of CD73 at the BBB in the tMCAO mouse model during the acute phase, indicating a fundamental role of this ectoenzyme in the inflammatory response to ischemic reperfusion in the brain. Our findings not only pave the way for future therapeutics targeting extracellular purinergic signaling but also highlight one of potentially multiple transcriptomic differences between mice and humans that may fundamentally impact disease response and the translation of preclinical findings.

## RESULTS

### Humanizing the ATP degradation machinery of murine brain endothelial cells

During ischemic stroke, brain cells release substantial amounts of the pro-inflammatory purine nucleotide ATP. To specifically assess the role of purinergic nucleotide conversion at the BBB, we first aimed to confirm the previously observed differential expression of key ectoenzymes between BECs of mice and humans. Subjecting primary brain microvascular EC from mice (mBMEC) and humans (hBMEC), as well as the murine BEC line bEnd.3, to qPCR analysis, we confirmed publicly available transcriptomic data[9, 18, 19]. Murine BECs predominantly expressed CD39, which converts extracellular ATP to ADP and AMP, whereas CD73, the enzyme that converts AMP to adenosine (ADO), was barely detectable. In contrast, human BEC predominantly expressed CD73, while the expression of CD39 was significantly lower (**Fig. 1A**). Assuming that the over 10-fold lower expression of CD73 in murine BEC compared to human BEC would fundamentally impact the BBB’s ability to generate anti-inflammatory ADO, which in turn might have brought implications on cerebral ischemic reperfusion injury, we generated an AAV vector mediating constitutive CD73 expression at the murine BBB, by employing the AAV-BR1 capsid, the ubiquitously active CAG hybrid promoter and woodchuck hepatitis virus posttranscriptional regulatory element (**Fig. 1B**). Transducing murine bEnd.3 BEC with the AAV-BR1-CAG-CD73 vector (“AAV-CD73”) yielded *Nt5e* transgene RNA expression levels (**Suppl. Fig. 1**), similar to the endogenous human *NT5E* levels (as shown in **Fig. 1A)**, and about 80% CD73-positive cells as assessed by flow cytometry, whereas CD73 could barely be detected in bEnd.3 cells treated with an empty control vector (**Fig. 1C**). Next, we assessed the capacity of murine and human BEC to degrade extracellular ATP to ADO and further breakdown products such as inosine (INO) by high-performance LC-MS/MS, to detect a potential shift from a pro-inflammatory to a more anti-inflammatory environment (**Fig. 1D**). 30 min after the addition of 100 µM ATP to hBMEC, we measured substantial levels of extracellular ADO (around 40 µM), while around 30 µM ATP were still found in the supernatant. Murine bEnd.3 cells, on the other hand, were able to break down the complete added ATP, presumably due to their high CD39 expression levels, but they accumulated high concentrations of extracellular AMP (53 µM) and only generated low amounts of ADO (5 µM) and further breakdown products due to the lack of CD73. Treatment of murine bEnd.3 cells with the AAV-CD73 vector, however, equipped them with the ability to further convert most of the ATP metabolite AMP into ADO (20 µM), INO (25 µM), and other purines. When bypassing CD39 and directly adding 100 µM AMP, hBMEC degraded almost 80% of the said AMP to ADO and INO, while murine bEnd.3 cells, conversely, only degraded about 20% of the added AMP into ADO and INO. Upon AAV-CD73 vector treatment, however, murine bEnd.3 cells converted about 75% of the added AMP to ADO (40 µM) and INO (20 µM) and further metabolites, attesting to an almost “humanized” degradation capacity of extracellular purinergic nucleotides by vector-transduced murine BECs (**Fig. 1D**). The purine metabolites hypoxanthine, xanthine, tryptophane, and nicotinamide (NAD) were either non-detectable or present at only very low concentrations (**Supplementary Figure 2**).

**Figure 1.**
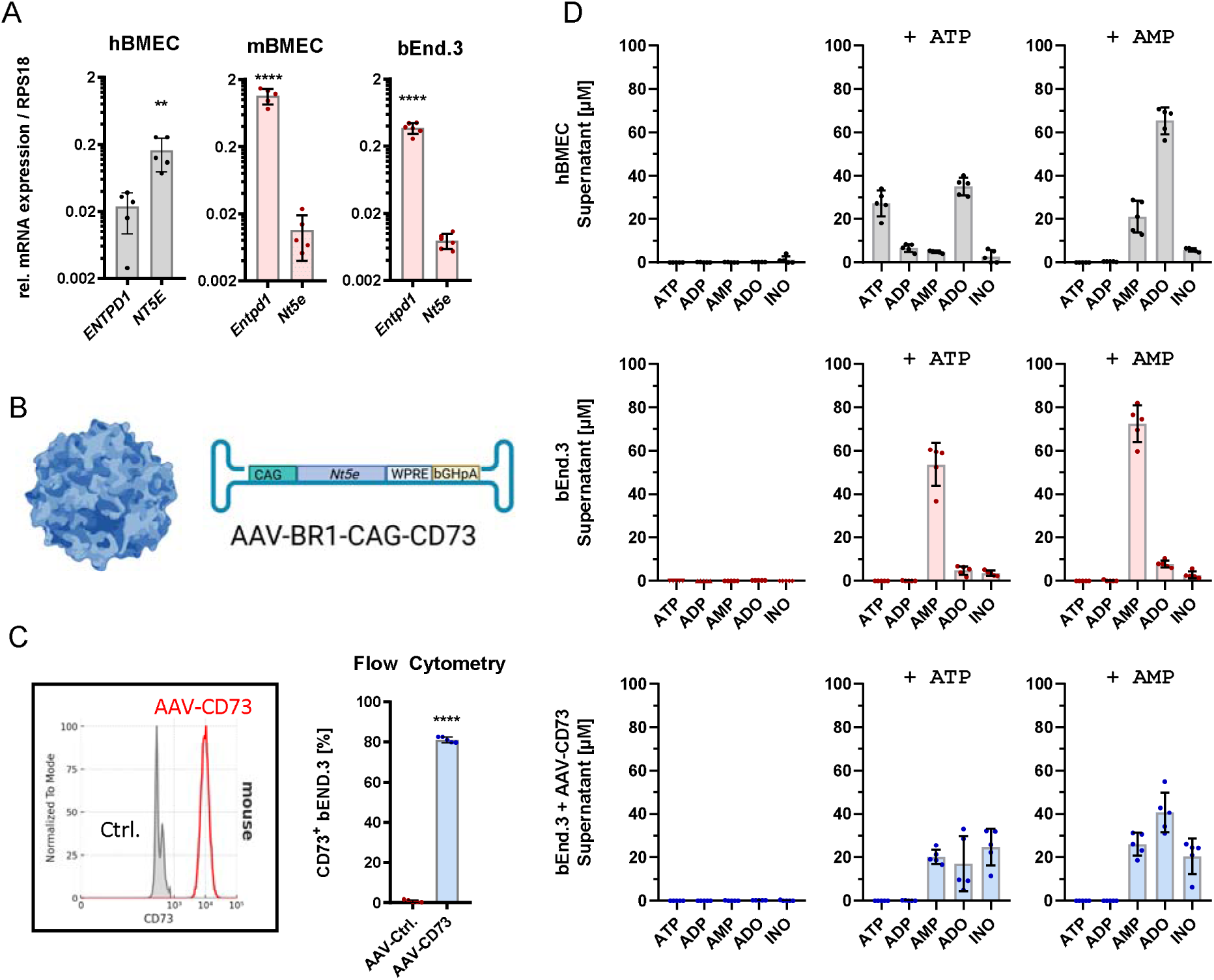
Species-dependent degradation of extracellular purinergic nucleotides by brain endothelial cells. **A**) Expression of *ENTPD1* (CD39) and *NT5E* (CD73) mRNA in brain endothelial cells from humans and mice relative to the RPS18 housekeeping gene. Primary human brain microvascular endothelial cells (hBMEC), primary murine brain microvascular endothelial cells (mBEMC), and the murine brain endothelial cell line bEnd.3 show inverse expression levels of *ENTPD1* and *NT5E* between humans and mice (n = 5 independent experiments/ cell type). Statistics: Two-tailed t-test; ** p = 0.0071; **** p <0.0001. **B**) Schematic overview of the AAV-BR1-CAG-CD73 vector used to “humanize” the expression level of *Nt5e* in murine brain endothelial cells. The brain endothelial cell-targeted AAV-BR1 capsid was employed to deliver the murine *Nt5e* (CD73) coding sequence under control of the strong ubiquitous CAG promoter and the bovine growth hormone polyadenylation signal (bGHpA), enhancing expression by the woodchuck hepatitis virus posttranscriptional regulatory element (WPRE). **C**) Transduction of murine brain endothelial cells (bEnd.3) by AAV-BR1-CAG-CD73 (100,000 genomic particles/cell) yields 80% CD73-positive cells as assessed by flow cytometry compared to cells transduced with an empty control vector being negative for CD73 (n = 5 independent experiments). Statistics: Two-tailed t-test; ****= p <0.0001. **D**) The concentration of extracellular purinergic nucleotides in the supernatant of primary human brain microvascular endothelial cells (hBMEC; upper panel), murine brain endothelial cells (bEnd.3; intermediate panel) and murine brain endothelial cells (bEnd.3) treated with AAV-BR1-CAG-CD73 vector (100,000 genomic particles/cell; lower panel) was determined by LC-MS/MS. Probes were collected 30 minutes after challenging the cells either with 100 µM ATP (middle) or 100 µM AMP (right) and compared to unchallenged cells (left). A total of n = 5 independent experiments were performed per cell type and treatment.

### Expression of CD73 in murine brain endothelial cells changes the transcriptional response to ATP

Having established that endothelial CD73 enhances extracellular ADO generation, we next investigated its effects on the transcriptional profile of brain endothelial cells. To mimic the release of extracellular ATP during ischemic stroke, two transgenic lines of bEnd3 cells were transiently exposed to 100 µM ATP: bEnd.3-CD39^ko^ cells lacking *Entpd1* and expressing low endogenous *Nt5e* levels, and bEnd.3-CD73^tg^ expressing high levels of CD39 and CD73. Interestingly, the capacity to degrade ATP to ADO substantially affected the brain endothelial transcriptomic response to extracellular ATP as assessed by Gene Set Enrichment Analyses (GSEA). bEnd.3-CD73^tg^ cells responded to ATP stimulation by enriching the Hallmark GO terms ‘cell junction organization’ (GO: 0034330) and ‘immune system process’ (GO: 0002376) (**Fig. 2A**), while genes of the same GO terms were downregulated in bEnd.3-CD39^ko^ cells (**Fig. 2B**), suggesting that increased ADO generation by brain endothelial cells might influence immune cell infiltration. Vice versa, GO terms indicating mitochondrial stress and enhanced cellular metabolism, such as ‘protein folding’ (GO: 0006457), ‘lyase activity’ (GO: 0016829), ‘mitochondrion’ (GO: 0005739), or ‘ribosome’ (GO: 0005840) were downregulated in bEnd.3-CD73^tg^ cells upon ATP treatment, presumably as consequence of rapidly converting ATP to ADO, while the same GO terms were upregulated upon ATP treatment in bEnd.3 CD39^ko^ cells (**Fig. 2A & B**). On the individual gene level, only a few candidates were differentially expressed above the threshold of fold change ≥ 2. Among these differentially expressed genes (DEGs), the important immunomodulator *Nfkbiz* was upregulated in ATP-treated bEnd.3-CD73^tg^ cells (**Fig. 2C**), which is in line with the detected induction of cell junction organization and immune system processes, whereas it was downregulated in bEnd.3 CD39^ko^ cells upon ATP treatment (**Fig 2D**). *Nfkbiz* has been shown to mediate neuroprotection and BBB integrity[20], therefore suggesting a protective effect of brain endothelial CD73 expression in the context of stroke.

**Figure 2.**
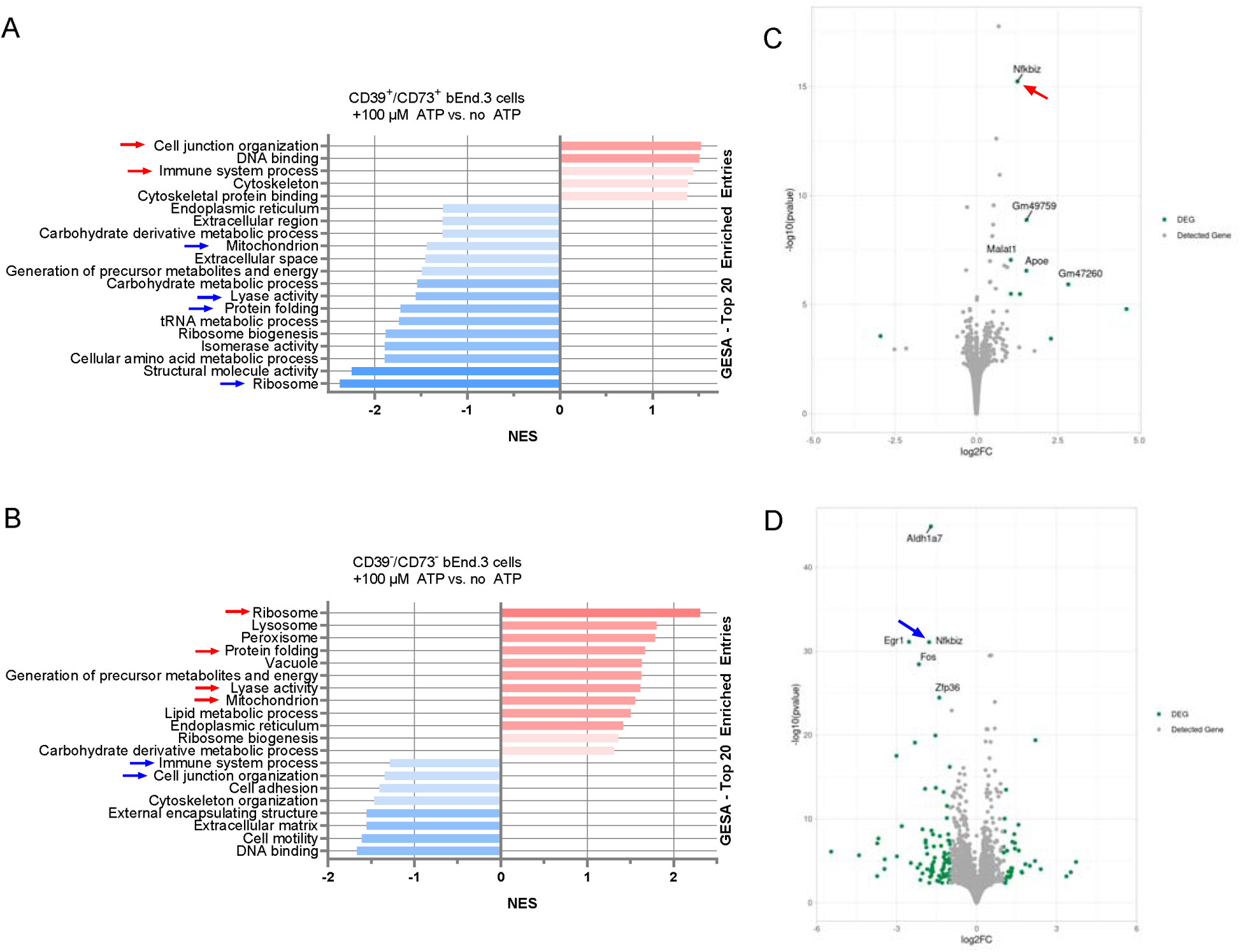
Ectoenzyme-dependent transcriptional response of brain endothelial cells to ATP. **A**) Gene set enrichment analysis (Hallmark) of ATP-treated (100 µM) bEnd.3-CD73^tg^ cells expressing high levels of *Entpd1* and *Nt5e* shows enrichment of ‘cell junction organization’- (GO: 0034330) and ‘immune system process’- (GO: 0002376) related genes, whereas GO terms indicating enhanced cellular metabolism and mitochondrial stress, such as ‘protein folding’ (GO: 0006457), ‘lyase activity’ (GO: 0016829), ‘mitochondrion’ (GO: 0005739), and ‘ribosome’ (GO: 0005840), were downregulated. **B**) Gene set enrichment analysis (Hallmark) of 100µM ATP-treated bEnd.3-CD39^ko^ cells lacking *Entpd1* and barely expressing *Nt5e* reveals an almost inverse gene regulation compared to ATP-treated BECs expressing high levels of *Entpd1* and *Nt5e* as shown in Fig. 2A. **C**) Volcano plot indicating DEGs upon ATP treatment (100 µM) of BECs expressing high levels of *Entpd1* and *Nt5e.* **D**) Volcano plot indicating DEGs upon ATP treatment (100 µM) of BECs expressing no *Entpd1* and low levels of *Nt5e*.

### Expression of CD73 at the murine BBB improves stroke outcome and prevents early mortality

After analyzing ADO generation by BECs and its consequences *in vitro*, we next validated the AAV-CD73 vector for its ability to mediate CD73 expression *in vivo*. AAV-CD73 was administered to mice at three different doses (between 2.5 x 10^11^ and 2.5 x 10^12^ vg/ animal) by tail vein injection, and CD73 expression was assessed 14 days later. As expected, we saw a dose-dependent increase in viral copy numbers in brain tissue (**Fig. 3A**), yielding a dose-dependent increase in transgenic *Nt5e* (CD73) expression, while the low expression level of endogenous *Nt5e* was not affected (**Fig. 3B**). Further, while BECs isolated from mice treated with an empty control vector were CD73-negative in flow cytometry analysis, AAV-CD73-treatment yielded 70-90% CD73-positive BECs, depending on the dose (**Fig. 3C**). Only very few AAV-BR1 vector copies were detected in the liver, an organ commonly targeted by many AAVs, attesting to the high specificity of the utilized viral vector for BECs (**Suppl. Fig 2**).

**Figure 3.**
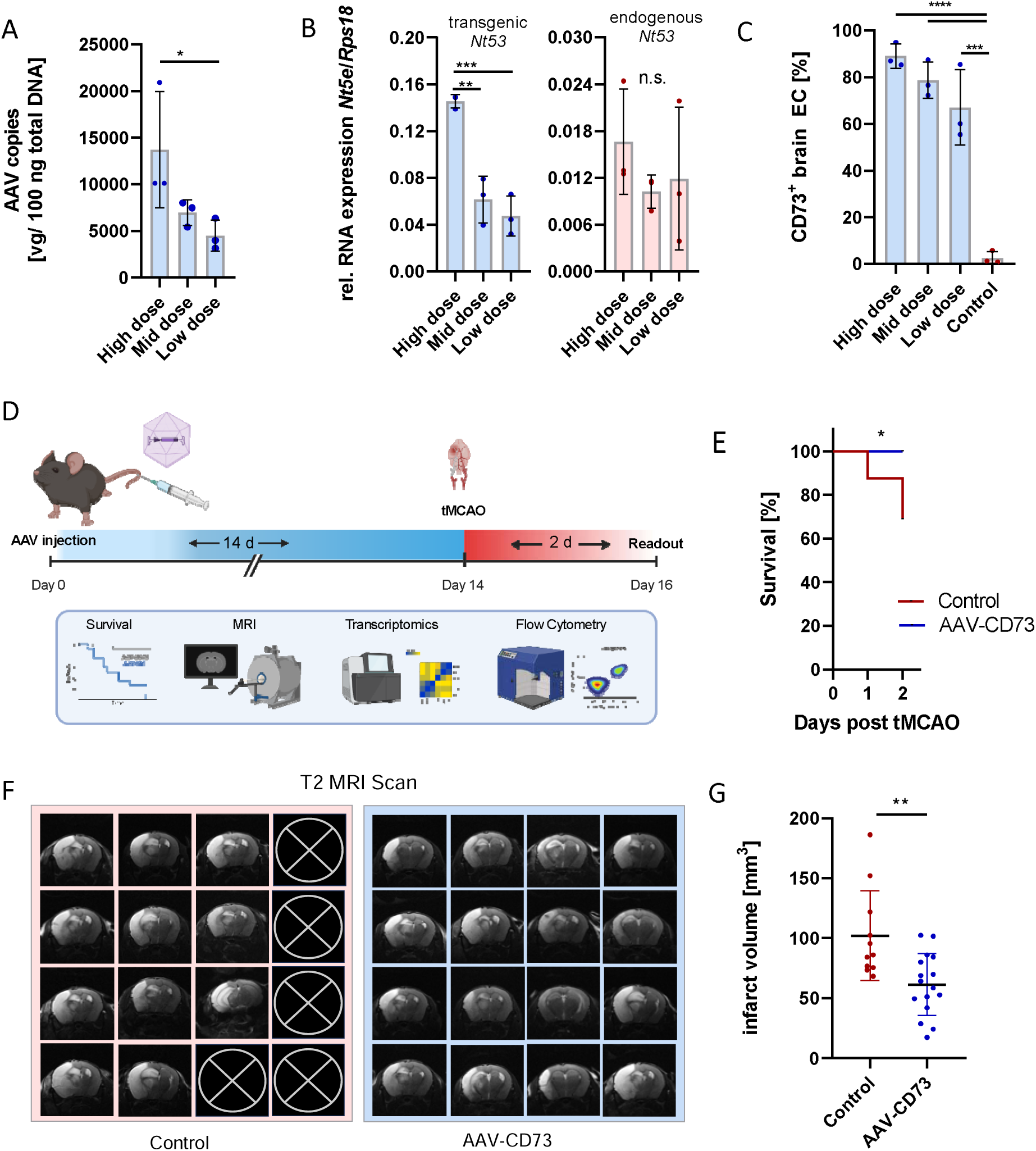
Expression of CD73 at the murine blood-brain barrier is protective in ischemic stroke. **A**) Dose finding experiments were conducted to validate the AAV-mediated expression of CD73 in murine brain endothelial cells *in vivo*. Mice either received 5 x 10^11^ (low dose), 1 x 10^12^ (mid dose), or 2.5 x 10^12^ (high dose) genomics particles (gp)/ animal via the tail vein. The number of AAV vector copies in the brains of n = 3 mice/ group was determined by qPCR 14 days after vector administration. Statistics: Kruskal-Wallis test; * p = 0.0328. **B**) The expression of Nt5e mRNA relative to RPS18 was determined by qPCR in n = 3 mice/ group 14 days after vector administration, distinguishing between transgenic (left) and endogenous *Nt5e* (right). Statistics: One-way ANOVA followed by Turkey’s multiple comparison; ** p = 0.0014; *** p = 0.0006; n.s. = not significant. **C**) CD73-positive brain endothelial cells isolated from AAV-BR1-CAG-CD73-treated mice were quantified by flow cytometry in comparison to cells isolated from mice treated with empty control vector 14 days after vector administration (n = 3 mice/ group). Statistics: One-way ANOVA followed by Turkey’s multiple comparison; *** p = 0.0001, **** p <0.0001. **D**) Schematic overview of the experimental setup used to assess the effect of brain endothelial CD73 on ischemic stroke. Mice received AAV-BR1-CAG-CD73 or empty control vector at a dose of 2.5 x 10^12^ gp/ animal via tail vein injection. 14 days after AAV administration, mice underwent transient middle artery occlusion (tMCAO). The effect on the stroke outcome was assessed in the late acute phase, 2 d after tMCAO by T2-weighted MRI (infarct volume), flow cytometry (cerebral immune cells composition), and bulk RNAseq of isolated BEC as well as of isolated brain leukocytes. The survival of all animals was monitored until the final experiments 2 d post-tMCAO. **E**) Kaplan-Meyer curve indicating the improved survival of mice expressing CD73 at the BBB. While all mice in the AAV-BR1-CD73-treated group survived until the final experiments at day 2 post tMCAO (n = 16), only 11 out of 16 mice survived until day 2 in the control group. Statistics: Log-rank (Mantel-Cox) test; * p = 0.0166. **F**) Infarct volume measured by T2-weighted MRI. Coronal MRI sections (section 14 out of 24) from each measured mouse (n = 11 mice/Ctrl; n = 16 mice/AAV-CD73). **G**) Total infarct volumes assessed by T2-weighted MRI. Statistics: Two-tailed t-test; ** p = 0.0025.

Next, we assessed the effect of purinergic nucleotide conversion at the BBB on ischemic stroke. We injected mice intravenously with the highest vector dose tested (2.5 x 10^12^ vg/animal), assuming that the strongest CD73 expression would yield the most efficient generation of extracellular ADO. As control, we used an AAV-BR1 vector carrying no transgene (“Ctrl.”) at the same dose. Fourteen days after vector application, mice underwent tMCAO surgery followed by a recovery phase of two days. Survival was monitored, and the outcome was assessed in the acute phase, 2 d after tMCAO surgery. A schematic overview of the experimental setup is shown in **Figure 3D**. AAV-CAG-CD73 treatment significantly (p = 0.0166) increased the survival (**Fig. 3E**). While 5 out of 16 mice in the control group died by day 2 (31.25%), all 16 mice in the AAV-CD73-treated group survived the same period (100%). In line, the infarct volume of animals in the AAV-CD73-treated group measured by T2-weighted MRI reduced by 40% compared to animals treated with the empty control vector (**Figs. 3F & G**).

### Improved CD73-induced stroke outcome correlates with a shift towards myeloid cells

As the GO term ‘immune system process’ was one of the few being upregulated by murine BECs expressing CD73 upon ATP treatment (**Fig. 2**), we hypothesized that the observed protective effect of endothelial CD73 expression on stroke outcome might be linked to immune cells. Thus, we next analyzed how brain endothelial CD73 expression affects leukocytes in the context of ischemic stroke *in vivo*. The impact of brain endothelial CD73 on leukocyte recruitment was assessed in the acute phase of stroke (d2) by flow cytometry, and exploratory transcriptomics (bulk RNA-Seq on cerebral CD45+ cells). A multi-channel gating strategy was employed to distinguish between nine types of leukocytes (**Figure 4A)**. Significant differences in the cerebral leukocyte counts between AAV-CD73-treated mice and controls were neither detected for microglia, monocytes/ macrophages, neutrophils, NK cells, γδ T-cells, nor classical CD4^+^ or CD8^+^ T-cells (**Fig 4B**). However, monocytes/ macrophages showed a significant relative enrichment in the brains of AAV-CD73-treated mice when normalizing the cell count to the infarct volume (**Fig. 4C)**. Relative to the infarct volume, we also observed significantly more γδ T-cells (**Fig 4D**), although they were least abundant among all analyzed cell types.

**Figure 4.**
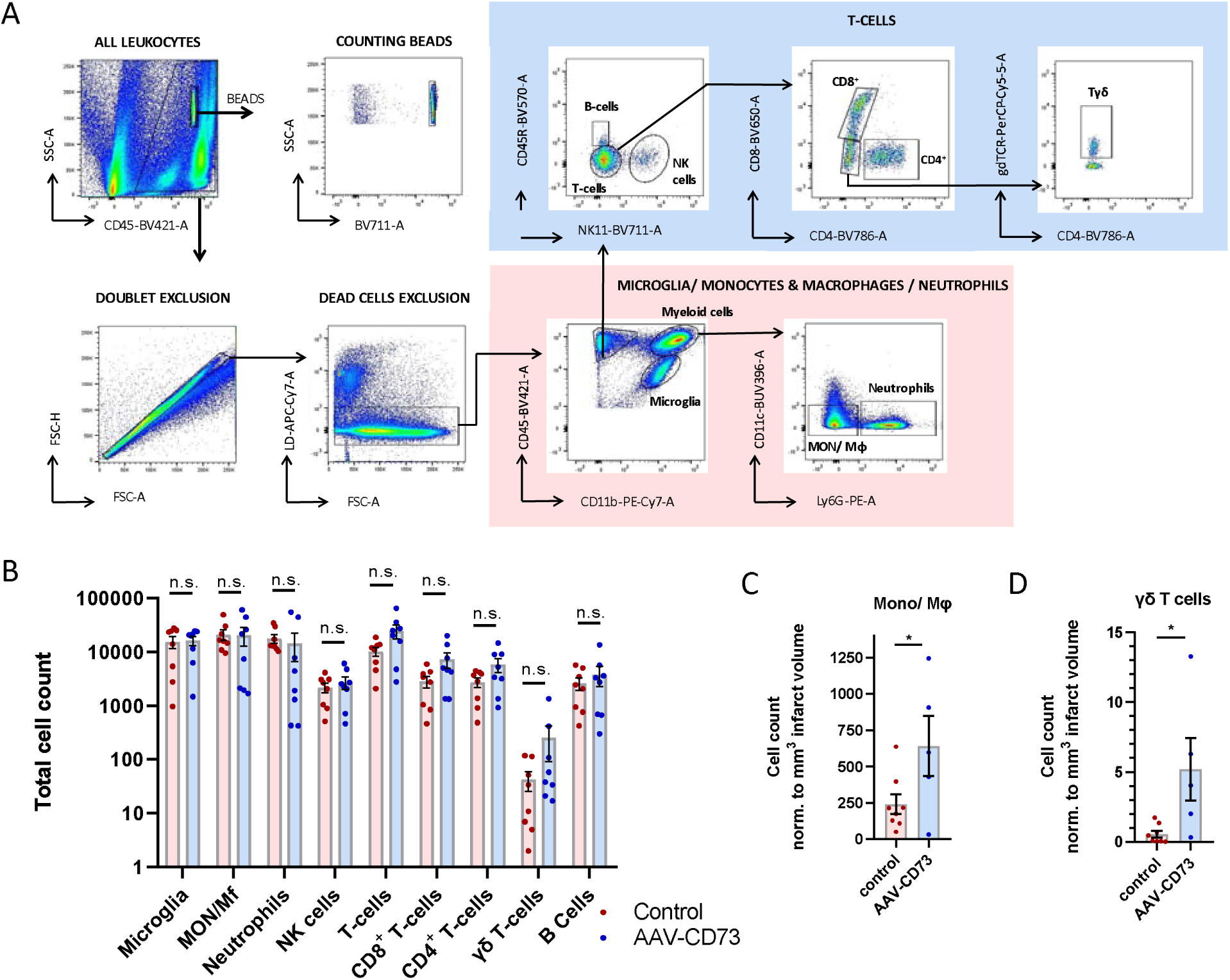
Endothelial CD73-induced brain leukocyte infiltration. The effect of brain endothelial CD73 expression on infiltrating leukocytes was assessed *in vivo* 16 d after intravenous AAV injection (2.5×10^12^ gp/animal) in the acute phase, 2 d after tMCAO. **A**) Flow cytometry gating strategy. Counting beads were used to normalize cell counts. Doublets and dead cells were excluded. CD45^+^/CD11b^-^ leukocytes were divided into CD45R^+^/NK1.1^−^ B cells, CD45R^−^/NK1.1^+^ NK cells. and CD45R^−^/NK1.1^−^ T cells. T cells were further distinguished into CD4^+^ T cells, CD8^+^ T cells, and γδTCR^+^ T cells. Among the CD11b^+^ cells, microglia were defined as CD45^low^/CD11b^+^/P2Y12^+^, while infiltrating myeloid cells were defined as CD45^high^/CD11b^+^. The latter population was separated into Ly6G^+^ neutrophils and Ly6G^−^ monocytes/macrophages. **B**) Quantification of immune cells in whole brain single cell suspensions by flow cytometry, normalized to mm^3^ infarct volume (n = 8 mice/Ctrl; n = 5 mice/AAV-CD73). Statistics: Microglia, monocytes/macrophages, neutrophils: Two-tailed t-test: * p = 0.0485; n.s. = not significant. NK cells, T cells, CD8+ T cells, CD4+ T cells, γδ T cells, B cells: Mann-Whitney test: * p = 0.0186; n.s. = not significant.

Bulk RNA-Seq of cerebral leukocytes revealed numerous differentially expressed genes (680), the majority of which (656) were upregulated (**Fig. 5A**). Gene set enrichment analysis for affected molecular functions (MF) revealed a broad upregulation of chemotaxis-related genes involved in chemokine/cytokine activity and binding (**Fig. 5B**), indicating increased leukocyte recruitment and activation upon brain endothelial CD73 expression. A heatmap showing the expression level of all DEGs belonging to the GO term ‘chemotaxis’ (GO: 0006935) can be found in **Supplementary Figure 3A**. Focusing on the biological processes (BP), we found migration and chemotaxis of neutrophils and granulocytes to be among the top 20 enriched terms (**Fig 5C**). Further, GSEA suggested enhanced production of IFN-γ, TNF, as well as of several interleukins, supporting enhanced leukocyte activation (**Fig 5C**).

**Figure 5.**
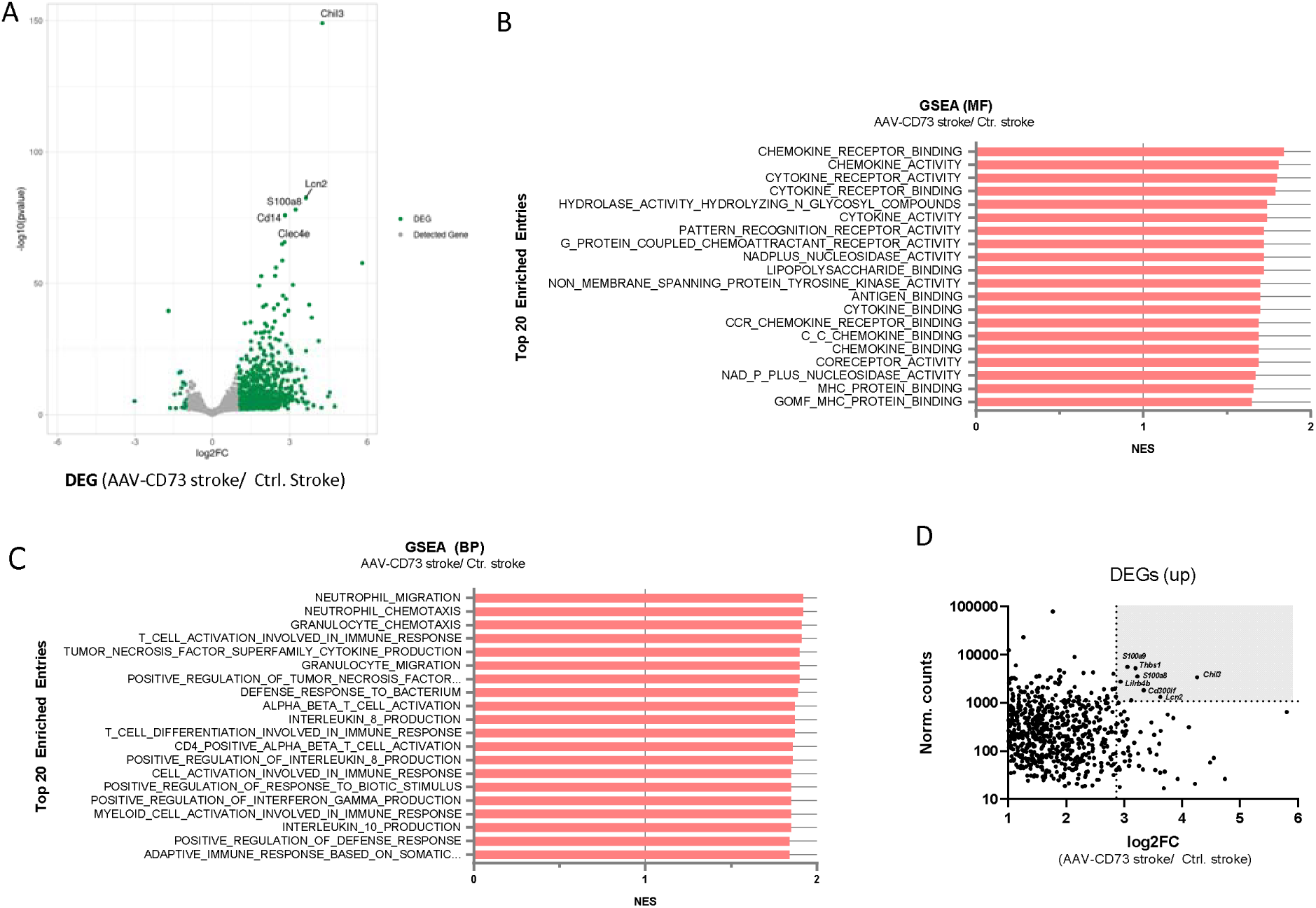
Expression of CD73 at the murine blood-brain barrier induces trancriptomic changes in brain leukocytes. **A)** Volcano plot indicating differentially expressed genes (DEGs) in CD45^+^ leukocytes isolated from brains of mice with brain endothelial CD73 overexpression (AAV-CD73) compared to controls that received an empty control vector (Ctrl), 2d after tMCAO. **B**) Gene set enrichment analysis (Molecular Function) of CD45^+^ leukocytes isolated from brains of mice with brain endothelial CD73 overexpression (AAV-CD73) compared to controls that received an empty control vector (Ctrl), 2d after tMCAO. **C**) Gene set enrichment analysis (Biological Process) comparing CD45^+^ leukocytes isolated from brains of mice with brain endothelial CD73 overexpression (AAV-CD73) compared to controls that received an empty control vector (Ctrl), 2d after tMCAO. **D)** Dot plot indicating the most abundant and most strongly enriched DEGs in CD45^+^ leukocytes isolated from brains of mice with brain endothelial CD73 overexpression (AAV-CD73) compared to controls that received an empty control vector (Ctrl), 2d after tMCAO.

Of note, in addition to IL-8, production of the anti-inflammatory IL-10 was among the most strongly enriched biological processes, suggesting a shift towards inflammation resolution. To get a rough idea of the most profound CD73-induced transcriptomic changes on the individual gene level, we focused on the most abundant DEGs showing the biggest difference in expression between mice treated with BR1-CAG-CD73 and controls (**Fig 5D**). Some of the top candidates, such as S100a8 & *S100a9* (calprotectin), *Chil3,* and *Lcn2* correspond to amplified innate effector signals, whereas others, such as *Thbs1*, an inducer of Il-10 production[21], as well as the myeloid cell-derived immune receptors *Lilrb4* [22] and *Cd300lf[23]*, again, might indicate a beginning shift towards inflammation resolution. In line with the GSEA data, all of the ten most strongly upregulated genes whose expression levels were at least moderate (>100 normalized counts; **Suppl. Fig 3B**) were primarily attributed to cells of the myeloid lineage, mostly macrophages/ monocytes and neutrophils, which suggests increased myeloid cell recruitment. Among the 40 most abundant differentially expressed genes (>2000 normalized counts, **Suppl. Fig. 3C**), the vast majority, again, was primarily attributed to myeloid cells, which is in line with the relative increase of these cells as observed by flow cytometry. Interestingly, *Chil3*, one of the most strongly expressed and top upregulated genes, is not only considered an innate effector gene but also serves as a classical marker for alternatively activated rodent macrophages of an “M2-like” phenotype[24], which, in concert with the upregulated genes *Thbs1, Lilrb4*, and *Cd300lf,* yet again, might indicate a shift towards inflammation resolution [25] [23, 26]. In line, we also observed an increase in *IL4Ra,* which is known to associate with the above-mentioned *Cd300lf[27]*, and, if expressed on macrophages, can induce tissue repair and inflammation resolution[28]. For comparison, transcriptomic analysis has also been performed on CD45^+^ cerebral leukocytes from healthy mice. However, lacking ischemia as an immunogenic trigger, the expression profile of cerebral immune cells from AAV-CD73-treated mice that did not undergo tMCAO, presumably mostly microglia, did not differ from animals treated with a control vector (data not shown).

## DISCUSSION

Ischemic stroke remains a leading cause of death and disability worldwide, with secondary implications such as neuroinflammation and BBB dysfunction critically shaping the outcome. Beyond the initial vascular occlusion, the release of damage-associated molecular patterns (DAMPs), particularly extracellular ATP, drives inflammatory cascades that exacerbate neuronal injury via receptors expressed on a multitude of brain-resident and immune cells[29]. ATP acts as a potent danger signal, activating P2X and P2Y receptors on microglia, astrocytes, and infiltrating immune cells, amplifying neuroinflammation[9, 12, 30]. Degradation of ATP to adenosine through ectonucleotidases such as CD39 and CD73 counterbalances these effects, as adenosine signaling via adenosine receptors is thought to exert predominantly anti-inflammatory, vasodilatory, and tissue-protective effects[14]. The exact mechanisms of action, however, are complex as adenosine can act on four different G-protein-coupled receptors (A_1_, A_2A_, A_2B_, A_3_), two of which (A_2A_, A_2B_) signal mainly via G_s_-protein-induced adenylyl cyclase activation, whereas the other two (A_1_, A_3_) inhibit adenylyl cyclase activity via G_i_-proteins[13].

Given the enormous importance of purinergic signaling in shaping (neuro-) inflammatory responses, it appears surprising that human BECs constitutively express CD73, whereas murine BECs only exhibit minimal CD73 levels[9]. Even far less pronounced differences in CD39/CD73 expression have been shown to substantially influence inflammatory disorders such as cardiac fibrosis, where male patients show a better outcome than female patients due to higher expression levels of these critical ectoenzymes[31]. Our findings demonstrate that endothelial CD73 expression significantly improves survival by reducing infarct volume within the acute phase, 2d post-tMCAO, despite relatively increased leukocyte infiltration. Although immune cell infiltration into the brain is commonly associated with worse outcomes in neuroinflammatory diseases and stroke, our findings highlight that the number of infiltrating immune cells is not a predictive measure per se and strongly depends on the cells’ phenotype and activation status. Flow cytometry indicated increased infiltration, especially of monocytes/macrophages in mice expressing CD73 at the BBB 2d after tMCAO. Yet infarct sizes were substantially smaller, and survival was improved. Transcriptomic analyses indicated a pronounced effect of endothelial CD73 expression on infiltrating immune cells, especially on myeloid cells. Despite a strong inflammatory signature, we hypothesize to have improved tissue protection and repair by promoting chemotaxis and shifting macrophages towards a more anti-inflammatory phenotype, although this remains to be proven by future studies. A strong chemotactic effect of adenosine on neutrophils via the A_1_ receptor and an anti-apoptotic effect via the A_2A_ receptor have been demonstrated in the past[32–35], and a similar mode of action may also account for monocytes/macrophages. Our findings align with previous reports that adenosine signaling promotes macrophage polarization toward anti-inflammatory M2 states [25, 36, 37] and dampens cytotoxic T-cell activation [53, 54]. Of note, not only extracellular adenosine but also extracellular inosine was enhanced by endothelial CD73 expression. Alike adenosine, inosine has been shown to exert an anti-inflammatory effect[38]. The observed protective effect in our study might, therefore, have been caused by either one of the two adenine nucleotides or both. Our findings suggest that brain endothelial CD73 expression promotes a neurovascular environment, reducing injury, shifting the balance from detrimental inflammation to protective signaling. The big interspecies difference in CD39/CD73 expression between mice and humans, therefore, has implications for stroke research. Particularly, it adds to the ongoing discussion of how good current mouse models can predict the clinical benefit of novel immunomodulatory substances[39, 40]. Recently, it has been shown in a landmark study that the blood leukocyte response during the acute phase of stroke substantially differs between humans and mice[41]. The differential expression of molecules involved in purinergic signaling, such as CD73, might contribute to this effect.

Previous studies have reported conflicting roles for CD73 in ischemic injury. In photothrombotic models, global CD73 deficiency aggravated infarct size and immune cell infiltration, reversible by exogenous 5′-nucleotidase supplementation[15]. Conversely, in tMCAO, global CD73 knockout did not alter infarct size or leukocyte infiltration at day 3 post-stroke[16]. These discrepancies likely reflect differences in stroke models. Photothrombosis induces endothelial cell injury, thrombus formation, and BBB disruption, resulting in vascular edema, whereas tMCAO incorporates reperfusion, resulting in a penumbra with delayed parenchymal cell death and secondary immune reactions[9]. Further, a global knockout of CD73 does not take into account the pronounced cell-type-dependent differences in the expression level of this important enzyme. Given the very low expression level of CD73 in murine BECs, it is not surprising that its knockout did not yield meaningful effects in the tMCAO model[16], whereas the present study indicates a substantial benefit of brain endothelial CD73 overexpression in the same stroke model. Our targeted approach provides direct evidence that BBB-localized CD73 exerts strong protective effects in an ischemia-reperfusion setting, strengthening the case for cell-type–specific strategies in translational research[14].

Our findings underscore the huge therapeutic potential of enhancing purinergic signaling at the BBB, e.g., by the BBB-targeted delivery of ectoenzymes such as CD39 or CD73 in human stroke patients. Interestingly, mice have recently been shown to profit from an endothelial-directed delivery of CD39 in the tMCAO model, although murine BECs naturally express substantial levels of this enzyme, while lacking CD73. The benefits of endothelial CD39 delivery reported by Lee et al.[42], therefore, are more likely explained by an improved degradation of extracellular ATP as opposed to increased adenosine. The results of our study show that CD73-based strategies could complement this approach by accelerating the essential step of AMP-to-adenosine conversion. Further, our study highlights the limitations of current mouse models by demonstrating that mice with “humanized” CD73 expression levels show a vastly different inflammatory profile and outcome at least in the tMCAO reperfusion model of stroke.

A limitation of our study is the focus on the acute phase of stroke (48 h), while intermediate and long-term effects remain unknown. It has already been shown that macrophages tend to show a protective ‘M2’-like phenotype at early stages of ischemic stroke, whereas they can shift towards an ‘M1’-like phenotype in the later phase[43], although this seems to depend on the exact type of macrophage, as border-associated macrophages seem to impair BBB integrity and induce neurological impairments already in the acute phase[44]. Whether the observed strongly beneficial effects of brain endothelial CD73 expression on stroke outcome will change over prolonged observation periods, therefore, remains to be analyzed in future studies. It has recently been shown that, after stroke, the transcriptome of blood-borne myeloid cells in the brain remains distinct from their counterparts in the blood[41]. We, however, have not investigated whether the observed transcriptomic changes in the CD45^+^ cell fraction from the brain differ from those in the blood. Further, the proposed switch in the macrophage activation status, as suggested by bulk RNA sequencing, needs to be verified on the individual cell level, e.g. by single-cell RNA sequencing or in-depth flow cytometry profiling. Another limitation is the use of only male mice, precluding assessment of sex-specific responses, which is a highly critical consideration given the strong evidence for sex differences in stroke pathophysiology[45, 46]. As pronounced sex differences have not only been described for human stroke patients but also for mice[47], this study was designed to focus on only one sex to reduce the statistical variance and thereby the required number of animals used. Age is another factor having been shown to influence the immune response to stroke[48, 49]. In our study, we only used young adult animals. Future studies are needed to investigate the effect of enhanced ectonucleotidases at the BBB of female and aged mice as well as stroke patients.

While our AAV-BR1 approach effectively demonstrates a prototype for endothelial CD73 enhancement in mice, a direct translation to clinical settings remains unrealistic, especially due to the time required for AAV-mediated gene expression. However, our study provides very important mechanistic insights for future pharmacological strategies of adenosine enhancement, including CD73/CD39 delivery, e.g., through nanobodies [12] or the use of adenosine receptor agonists, and thereby proposes a promising direction for future investigations. While humans endogenously express substantial levels of CD73 at their BBB, human BECs express much less CD39 than mice. This renders pharmacologic enhancement of CD39 rather than CD73 at the BBB a promising treatment option for stroke patients to reduce extracellular ATP and increase adenosine.

## METHODS

### Cell Culture

Murine brain endothelial cells (bEnd.3; ATCC #CRL-2299) and human embryonic kidney 293T cells (HEK 293T; DSMZ #ACC 635) were cultured in DMEM (Gibco) supplemented with 10% fetal bovine serum (FBS) and 1% penicillin-streptomycin at 37⍰°C and 5% CO₂. *Spodoptera frugiperda* cells (Sf9; Gibco #11496015) were maintained in serum-free Insect-XPRESS™ (Lonza) containing 10 mg/l gentamicin at 27 °C. Primary human brain microvascular endothelial cells (HBMEC; ScienCell, #1000) tested negative for HIV, HBV, and HCV) were cultured in Endothelial Cell Medium (ECM) consisting of 500 ml basal medium (ScienCell), 25 ml FBS, 5 ml Endothelial Cell Growth Supplement (ScienCell) and 5 ml of penicillin/streptomycin solution and (ScienCell) and the culture flasks were coated with Fibronectin (Sigma-Aldrich). HBMEC were used for experiments not later than passage 5. All cells were routinely tested for mycoplasma contamination using the MycoCheck Mycoplasma PCR Detection Kit (ABP Biosciences, #A058-2).

### Assessment of brain endothelial CD73 expression levels

RNA was isolated from primary human (hBMEC) and murine (mBMEC) brain endothelial cells, the murine brain endothelial cell line bEnd.3 and murine brains using the Quick-RNA Miniprep Kit (Zymo Research). Nucleic acid concentrations and purity were assessed by NanoDrop spectrophotometry. RNA was used for cDNA synthesis using the High-Capacity cDNA RT-Kit (Applied Biosystems). Expression of human Nt5e (CD73) (Forward Primer: CCAGTACCAGGGCACTATCTG, Reverse primer: TGGCTCGATCAGTCCTTCCA), endogenous murine Nt5e (Forward Primer: CCTGCACACAAACGACGTG, Reverse primer: CTGGTCTCCGGCATCCAAAA) or transgenic murine Nt5e (Forward Primer: GAAGGAGGAACCAAACGTGCT, Reverse Primer: GGTGTACAGGAAGGTATTTGAG), was compared to human RPS18 (Forward Primer: GCGGCGGAAAATAGCCTTTG, Reverse primer: GATCACACGTTCCACCTCATC) and mouse RPS18 (Forward Primer: GCTCGCGGG TTGAGAGAAG, Reverse Primer: ACTGAGCAACAACACATCCTC) by the delta CT method.

### AAV vector plasmids

To generate the AAV transfer plasmid, a codon-optimized sequence for murine CD73 (ecto-5′-nucleotidase, *Nt5e*) was synthesized (GenScript) and cloned under the control of the CAG promoter (CMV early enhancer/chicken β-actin promoter) into a modified pFastBac1 plasmid (Thermo Fisher Scientific, #10359-016) containing the woodchuck hepatitis virus posttranscriptional regulatory element (WPRE) and the bovine growth hormone poly A signal sequence (bGHpA) flanked by AAV2 inverted terminal repeats (pFB1-CAG-CD73-WPRE). As control, we used the insert-less counterpart (pFB-CAG-empty-WPRE). As packaging plasmid, we used pFBD-AAV-BR1-Rep/Cap containing an AAV2 rep gene and a BEC-specific AAV-BR1 cap gene cloned into the pFASTBAC™ Dual vector (Thermo Fisher Scientific, #10712-024)

### Recombinant AAV production

Recombinant bacmid DNA was generated using the Bac-to-Bac™ Baculovirus Expression System (Thermo Fisher Scientific, #10359-016). The above-mentioned pFastBac plasmids were transformed into DH10Bac competent *E. coli* cells, which include the baculovirus shuttle vector (bacmid) and a helper plasmid encoding the Tn7 transposase. Bacmid DNA was isolated from transformed bacteria by alkaline lysis and ethanol precipitation. Recombinant baculovirus stocks encoding the AAV packaging functions (AAV2 Rep and AAV-BR1 Cap) and the CD73 vector genome or the empty control were generated by transfecting *Spodoptera frugiperda* Sf9 cells with the above-mentioned plasmids using TransIT Transfection Reagent (Mirus Bio). Sf9 cells were maintained in serum-free Insect-XPRESS™ (Lonza) containing 10 mg/l gentamicin at 27 °C for 72 h. High-titer baculovirus stocks were harvested after 72 h and used to co-infect Sf9 cells for AAV production. After incubating Sf9 cells with the respective baculovirus stocks for 72 h, AAVs were harvested from cells and cell culture medium. 72 h post-infection under gentle agitation (110 rpm), cells were harvested by low-speed centrifugation (150 ×g, 5 min) and the cell pellets were resuspended in PBS-MK buffer. To release AAV particles, the cell suspension was subjected to three freeze-thaw cycles (alternating between –80 °C and 37 °C). AAV particles contained in the Sf9 cell culture medium were precipitated by mixing the cell-free medium with 10 g PEG-8000 and 5.8 g NaCl per 100 ml of medium and incubating for at least 24 h at 4°C, followed by 30 min centrifugation at 10,000 x g. The AAV-containing pellet was resuspended in PBS-MK. The AAV containing probes were further subjected to benzonase (MilliporeSigma) treatment (50 U/mL) for 30 min at 37 °C to degrade cellular and baculoviral DNA. The lysates were clarified by centrifugation (10,000 × g, 15 min), and the supernatant containing crude AAV-BR1-CAG-CD73-WPRE or empty control vectors was collected for purification by ultracentrifugation (see below).

### Purification of recombinant AAV vectors

Recombinant AAV particles were purified from the clarified Sf9 cell lysate by iodixanol gradient ultracentrifugation. The lysate was transferred to an ultracentrifuge tube and underlaid with a discontinuous iodixanol gradient consisting of 15%, 25%, 40%, and 54% (w/v) iodixanol layers prepared in OptiPrep solution (Sigma-Aldrich) in PBS-MK. Ultracentrifugation was performed at 58,000 × rpm for 70 min at 18 °C in a 70.1 Ti rotor (Beckman Coulter). After centrifugation, the opaque band at the interface of the 40% iodixanol layer (containing the enriched AAV particles) was carefully extracted using an 18-gauge needle. The rAAV-containing fraction was transferred to Amicon Ultra 100K molecular-weight cut-off centrifugal filter units (Millipore) for buffer exchange, and stored in PBS at −80°c until further use.

### Quantification of recombinant AAV by qPCR

Vector genome (vg) titers of the purified recombinant AAV vectors were determined by quantitative real-time PCR (qPCR). A pair of primers targeting the CAG promoter region of the vector genome was used to amplify a 190 bp fragment (Forward Primer: AACGCCAATAGGGACTTTC, Reverse primer: GTAGGAAAGTCCCATAAGGTCA). Reactions were set up using SYBR Green master mix (Roche) using the LightCycler® 96 instrument (Roche). Standard curves were generated using a ten-fold dilution series of a plasmid containing the CAG expression cassette of known copy number. Genome titers were calculated by comparing the Cq values of samples to the plasmid standard curve and reported as vector genomes per milliliter (vg/mL). For the quantification of vector genome copies in genomic DNA from murine tissue DNA samples were diluted to 10 ng/µL and 50 ng total DNA was used per reaction. Vector genome copies were normalized to input DNA and reported as vg per 100 ng genomic DNA.

### In vitro assessment of AAV-mediated CD73 gene transfer

To assess the efficacy of the AAV-mediated CD73 gene transfer in vitro, bEnd.3 cells were seeded in 12-well plates at a density of 2.5 × 10⁵ cells/well. After 24⍰h, cells were transduced with recombinant AAV at a multiplicity of infection (MOI) of 1 × 10⁵ vg/ cell. Recombinant AAV was diluted in basal medium and incubated with cells for 2⍰h, followed by the addition of full medium to a total volume of 2⍰mL. Expression of *Nt5e* (CD73) and the capacity to degrade extracellular adenine nucleotides were assessed after 24 - 72⍰h post-transduction. To assess surface CD73 expression by flow cytometry, cells were dissociated using TrypLE (Thermo Fisher Scientific), neutralized with complete medium, and spun down by centrifugation (170 × g, 5⍰min). Cells were washed twice with FACS buffer (PBS with 0.5% BSA and 0.5⍰mM EDTA), then incubated for 30–60⍰min at 4⍰°C with anti-mouse CD73 antibody (FITC-labeled; **see Table 1**). Washed samples were resuspended in 300⍰µL FACS buffer and analyzed on a LSRFortessa flow cytometer (BD Biosciences) At least 10,000 events were recorded per sample. Non-transduced and isotype-stained controls were included. To evaluate CD73 enzymatic activity in transduced cells, AMP degradation was quantified by high-performance liquid chromatography (HPLC). Non-transduced bEND-3 cells and HBMEC were used as controls. Cells (2.5 × 10⁴ cells/well) were grown in serum-free medium to eliminate confounding ATPase/AMPase activity from serum. Following transduction, cells were incubated with either extracellular AMP or ATP (100⍰µM final concentration) for 30 minutes at 37⍰°C in serum-free medium. Supernatants were collected, centrifuged to remove debris, and immediately stored at –80⍰°C until analysis.

**Table 1.** List of used antibodies.

| Antigen | Clone | Fluorochrome | Supplier | Catalog Number | Target Cell Population | Dilution |
| --- | --- | --- | --- | --- | --- | --- |
| **CD45** | 30-F11 | PerCP-Cy5.5 | BioLegend | 103132 | All leukocytes | 1:100 |
| **CD11b** | M1/70 | BUV737 | BD | 612800 | Myeloid cells | 1:100 |
|  |  |  | Bioscience<br>s |  |  |  |
| **Ly6G** | 1A8 | AF700 | BioLegend | 127622 | Neutrophils | 1:100 |
| **P2Y12** | S16007D | APC | BioLegend | Custom | Microglia | 1:100 |
| **CD73** | TY/11.8 | FITC | BioLegend | 127219 | Ecto-enzyme<br>marker | 1:100 |
| **CD73** | AD2 | APC | BioLegend | 127210 | Ecto-enzyme<br>marker (alt.) | 1:100 |
| **CD4** | RM4-5 | BV785 | BioLegend | 100547 | T helper cells | 1:100 |
| **CD8** | 53-6.7 | BV650 | BD<br>Bioscience<br>s | 563786 | Cytotoxic T cells | 1:100 |
| **NK1.1** | PK136 | BV710 | BioLegend | 108745 | NK cells | 1:100 |
| **CD11c** | HL3 | BUV395 | BD<br>Bioscience<br>s | 564080 | Dendritic cells | 1:100 |
| **F4/80** | BM8 | BV605 | BioLegend | 123129 | Macrophages | 1:100 |
| **B220** | RA3-6B2 | BUV805 | BD<br>Bioscience<br>s | 748867 | B cells | 1:100 |
| **TCR$\gamma\delta$** | GL-3 | PerCP-Cy5.5 | BioLegend | Custom | $\gamma\delta$ T cells | 1:100 |
| **MHC II** | M5/114.15.<br>2 | FITC | BioLegend | 107606 | Antigen-<br>presenting cells | 1:100 |
| **CD39** | DuHa59 | APC | BioLegend | 143810 | Ectonucleotidas<br>e marker | 1:100 |
| **CD31** | MEC13.3 | BUV395 | BD<br>Bioscience<br>s | 740231 | Endothelial cells | 1:100 |
| **Live/Dead**<br>**dye** | — | Near-IR | Invitrogen | L10119 | Dead cell<br>exclusion | 1:1000 |

### Liquid chromatography–tandem mass spectrometry (LC-MS/MS)-based targeted metabolomics

Metabolites of the ATP degradation pathway were quantified in cell culture supernatants (see above) by high-performance LC-MS/MS as previously described[50]. Supernatants were thawed on ice, aliquoted in ice-cooled 1.5 ml reaction tubes (Eppendorf), and spiked with ^13^C_6_-nicotinamide (Sigma-Aldrich), as an internal standard. Adenine nucleotides were extracted by adding pre-cooled acetonitrile/water 8:2 (v/v) to the sample and vortexing thoroughly at highest level for 1 min. Following homogenization, samples were incubated for 10 min on ice and centrifuged at 4°C and 16,000xg for 10 min. The supernatant was transferred to a brown glass vial (Phenomenex) and tightly closed with a crimped cap (LABSOLUTE®). The extracts were measured immediately after extraction using a triple quadrupole mass spectrometer (QTRAP® 5500; SCIEX) coupled to an ultra-high-pressure liquid chromatography system (Nexera X2; Shimadzu). Analytes were separated by hydrophilic interaction liquid chromatography using a Luna® HPLC column (3 µm NH2 100 Å 150mm x 2 mm, Phenomenex). The autosampler was cooled to 4°C, column oven temperature was set to 25°C and injection volume was 10 µl. Solvent A contained 20 mM ammonium acetate in MS-grade water (pH 9.8) and solvent B contained 100% acetonitrile. The flow rate was set to 0.25 mL/min with a linear gradient of decreasing solvent B (0-17 min, 80%-0%), followed by 8 min at 0% solvent B and 5 min re-equilibration at 80% solvent B. The MS was equipped with an ESI source and all measurements were carried out in negative ionization and multiple reaction monitoring (MRM) scan mode. Data analysis was performed using SCIEX OS software (version 2.0.0.45330) and quantification in Microsoft Excel. Analyte peaks were identified based on MRM transitions and retention time. A standard calibration curve ranging from 2 nM to 2 µM was used for accurate quantification and all analyte peak areas were normalized against internal standard peak areas. Extracellular nucleotide concentrations in the supernatant of cells were corrected for endogenous metabolite levels in the respective cell culture media not containing any cells by substraction.

### Transcriptomics from murine brain endothelial cells

Murine bEnd.3 cell stably expressing *Nt5e* (CD73) or lacking *Entpd1* (CD39) were generated using the LeGO lentiviral system[51]. In brief, bEnd.3 cells stably expressing *Nt5e* were generated by transfecting cells with a lentiviral LeGO expression vector containing the murine *Nt5e* coding sequence. Murine bEnd.3 cells lacking *Entpd1*, were generated by transfecting cells with a lentiviral expression vector containing the coding sequence of Cas9 (LeGO-iG2/Neo-opt), in addition to a plasmid encoding murine *Entpd1*-directed guide RNA identified by the CHOPCHOP browser tool (http://chopchop.cbu.uib.no/). The genetically modified cells and wild-type bEnd.3 cells were seeded into six-well plates and grown in RPMI medium (Gibco) supplemented with 10% fetal bovine serum (FBS) and 1% penicillin-streptomycin at 37⍰°C and 5% CO₂. One day after seeding the cells, the medium was replaced by fresh medium either containing 100 µM ATP or not. 24 hours after the addition of ATP, the medium was removed, cells were washed with PBS, and snap-frozen immediately. Samples were stored at −80°C until further processing. RNA was isolated using the RNeasy Kit (Qiagen) according to the manufacturer’s instructions. RNA quality was assessed using the Bioanalyzer (Agilent). RNA-seq libraries were prepared by the GS Integrative Genomics Core Unit (NIG), University Medical Center Göttingen (UMG), Germany. Single-stranded DNA libraries were generated and sequenced on the Illumina HiSeq4000 platform with 50 bp reads.

### Animals

Wild-type C57BL/6J mice (12–16 weeks old) were used for all experiments. Animals were ordered through the animal facility of the University Medical Center Hamburg-Eppendorf from Charles River Laboratories. As the tMCAO model shows profound sex differences in infarct size and inflammatory response, only male animals were used throughout the study to reduce the variability, thereby decreasing group sizes. Animals were housed in individually ventilated cages with standard chow and water provided ad libitum. All mice were acclimatized to the animal facility for at least two weeks before experimental procedures. All procedures involving animals were approved by the responsible authority (Behörde für Justiz und Verbraucherschutz der Freien und Hansestadt Hamburg; Lebensmittelsicherheit und Veterinärwesen, Hamburg, Germany) and conducted in accordance with national and European laws as well as the ARRIVE guidelines.

### AAV-mediated CD73 delivery

To modulate CD73 expression in a cell-type-specific manner, mice received tail vein injections of AAV-BR1-CAG-CD73-WPRE or empty control vector at doses between 2.5 x 10^11^ and 2.5 x 10^12^ vg/animal in a total volume of 120 µL PBS, under brief inhalation anesthesia (2–2.5% isoflurane in oxygen). Vectors were administered 14 days before tMCAO to allow adequate transgene expression. Mice were monitored daily for well-being, health status, and behavior according to an approved scoring table until the end of the experiment.

### Dose-response testing of AAV-mediated CD73 delivery *in vivo*

To modulate CD73 expression in a cell-type-specific manner, mice received tail vein injections of AAV-BR1-CAG-CD73-WPRE or empty control vector at doses between 2.5 x 10^11^ and 2.5 x 10^12^ vg/animal in a total volume of 100 µL PBS, under brief inhalation anesthesia (2–2.5% isoflurane in oxygen). Mice were monitored daily for health status and behavior according to an approved scoring table until the end of the experiment. At 14 post injection, animals were sacrificed under deep isoflurane anesthesia by cervical dislocation. Whole brains were extracted, and both hemispheres were processed for flow cytometry while the cerebellum was snap-frozen in liquid nitrogen for nucleic acid isolation. RNA was isolated using the AllPrep DNA/RNA Micro Kit (Qiagen) following the manufacturer’s protocol, while DNA was isolated using the DNeasy Blood & Tissue Kit (Qiagen). Nucleic acid concentrations and purity were assessed by NanoDrop spectrophotometry. RNA was used for cDNA synthesis and qPCR analysis of transgene expression (see above), while genomic DNA was used for vector copy number quantification (see above). To specifically assess brain endothelial CD73 expression, brain hemispheres were processed using a microvessel enrichment protocol[16]. In brief, tissues were homogenized and centrifuged in 15% dextran to isolate the microvascular pellet. The resulting fraction was enzymatically digested and filtered through a 70⍰μm mesh. Single-cell suspensions were stained with a live/dead viability dye, anti-CD31, and anti-CD73 antibodies (**Table 1**). Flow cytometry was performed on a BD FACSymphony A3 cytometer, and data were analyzed using FlowJo software.

### Transient Middle Cerebral Artery Occlusion (tMCAO)

Age-matched mice were allocated randomly into treatment and control groups and coded by an independent researcher to achieve blinding of the experiments. Focal cerebral ischemia was induced via the filament-based tMCAO model. Mice were anesthetized with isoflurane (5% for induction; 1–2% for maintenance) and kept on a heating pad to maintain body temperature. The common, external, and internal carotid arteries were exposed. A silicon-coated monofilament (0.23 mm tip diameter) was inserted through the external carotid artery and advanced to occlude the middle cerebral artery (MCA). Cerebral blood flow was assessed by Laser-Doppler flowmetry, to ensure the desired ≥80% reduction in the ipsilateral vs. contralateral hemisphere. After 50 min, the filament was withdrawn to allow reperfusion. After wound sealing and disinfection, animals were placed back into their cages, which were kept on a heating pad for an additional 24 h. Animals received subcutaneous injections of pre-warmed Sterofundin® (0.5 mL) intraoperatively and postoperatively to prevent dehydration. Analgesia was provided with Buprenorphine (0.1 mg/kg s.c.) and Tramadol in drinking water (1 mg/mL) beginning 24 prior to surgery. Mice were monitored twice on the day of surgery and once each following day for well-being, health status, and behavior according to an approved scoring table until the end of the experiment. Exclusion criteria: Since successful tMCAO was assessed already during the surgery by Laser-Doppler flowmetry, only mice that did not survive 48 h post-tMCAO (the timepoint of further analysis) were excluded from analysis of infarct volume and immune cell infiltration.

### Magnetic Resonance Imaging (MRI)

MRI scans were performed in the acute phase after stroke, on day 2 post-ischemia using a 7T ClinScan MRI system. Mice were anesthetized with isoflurane (1.5–2%) and placed in a prone position on a heated bed with continuous monitoring of respiration, ECG, and body temperature. T2-weighted images (24 image scans per mouse) were acquired before and after intraperitoneal administration of gadolinium-based contrast agent (MultiHance®, 0.1 mmol/kg body weight). Infarct volumes were quantified from all 24 MRI scans per mouse by a blinded investigator using Fiji ImageJ. Immediately following MRI, the murine brains were processed for further analyses (see below).

### Assessment of immune cells in the brain by flow cytometry

Immediately following MRI, 2d after tMCAO, mice were euthanized via cervical dislocation under deep isoflurane anesthesia. After removing the cerebellum for downstream analysis, the remaining brain tissue was processed for flow cytometry. Brain hemispheres were enzymatically digested and processed following a modified protocol[16]. Briefly, tissue was incubated in digestion solution (DMEM with 1 mg/mL collagenase and 0.1 mg/mL DNase I) at 37⍰°C for 30 minutes. Following mechanical dissociation and Percoll gradient centrifugation, erythrocyte lysis was performed. After washing and resuspension in FACS buffer, cells were stained with a comprehensive antibody panel (CD45, CD11b, Ly6G, P2Y12, CD4, CD8, NK1.1; **Table 1**). Samples were analyzed using a BD FACSymphony A3 cytometer, and data were analyzed using FlowJo software by a blinded investigator. The following gating strategy was used: singlets were selected by FSC-A versus FSC-H, and dead cells were excluded using the live/dead viability dye. CD45^+^ leukocytes were identified after removal of beads and debris. Within the CD45^+^ population, B cells (CD45R^+NK1.1^−), NK cells (CD45R^−^/NK1.1^+^), and T cells (CD45R^−^/NK1.1^−^) were distinguished. T cells were further subdivided into CD4^+^ and CD8^+^ subsets, with γδ T cells identified as γδTCR^+^ within the T-cell compartment. Among CD11b^+^ cells, microglia were defined as CD45^low^/CD11b^+^/P2Y12^+^, while infiltrating myeloid cells were defined as CD45^high^/CD11b^+^. The latter population was separated into Ly6G^+^ neutrophils and Ly6G^−^ monocytes/macrophages. Cell counts were normalized to counting beads and mm3 infarct volume. A total of n = 8 mice per group were planned to be analyzed, of which n = 3 mice died within the control group before the experiment.

### Transcriptomics from isolated cerebral immune cells

Immediately following MRI, on day 2 after tMCAO, mice were euthanized via cervical dislocation under deep isoflurane anesthesia. The cerebellum was removed and snap-frozen for downstream analysis. The remaining brain tissue was processed for transcriptomics. Tissue dissociation and subsequent processing followed the protocols described above. To isolate immune cells, brain hemispheres were enzymatically digested and processed[16], stained with the above-mentioned MACS beads and sorted for CD31^-^, CD45^+^ cells. RNA was isolated using the Quick-RNA Miniprep Kit (Zymo Research) according to the manufacturer’s instructions. RNA quality was assessed using the Bioanalyzer (Agilent); only samples with RNA integrity number (RIN) ≥7 were processed for sequencing. RNA-seq libraries were prepared by BGI Genomics (Shenzhen, China) using the RNAref transcriptome service. Poly(A) mRNA was enriched using oligo(dT) magnetic beads and fragmented. First- and second-strand cDNA synthesis was performed using random N6 primers, followed by end repair, adapter ligation, and PCR amplification. Single-stranded circular DNA libraries were generated and sequenced on the DNBSEQ platform with paired-end 150 bp reads (PE150). Each sample yielded an average of ∼6.84 Gb of high-quality sequencing data.

### Bioinformatics Analysis

Sequence reads were processed using fastp v0.23.2 [52] to remove adapter-derived sequences and low quality bases, applying the software’s default parameters. Reads were then aligned to the mouse reference assembly (GRCm39.110) using STAR v2.7.10a[53]. Differential expression was assessed using DESeq2 v1.34.0[54]. Significant differential expression was defined by an absolute log₂ fold change of at least 1 combined with a false discovery rate (FDR) threshold of 0.1. Overrepresentation (ORA) and gene set enrichment analysis (GSEA) were performed with clusterProfiler v3.14 in combination with a selection of the Molecular Signatures Database (MSigDB) subsets C2 and H, i.e. Biocarta KEGG, Reactome, WikiPathways, the Hallmark gene sets, and Gene Ontology (GO). Transcription factor activity predictions are based on DoRothEA (v1.6.0)[55]. The BEC (bEnd.3) dataset contained 29 samples across six experimental groups: CD39ko_plusATP (n = 6), CD39ko_minusATP (n = 5), CD73_plusATP (n = 5), CD73_minusATP (n = 5), WT_plusATP (n = 4), WT_minusATP (n = 4). The leukocyte dataset originally included twelve samples across four experimental groups: CD73_no_stroke (n = 3), CD73_stroke (n = 3), Control_no_stroke (n = 3) and Control_stroke (n = 3) of which only n = 2 samples per condition were kept for the final analysis due to fundamental batch differences as assessed by PCA analysis.

### Blinding

The investigators performing the tMCAO surgery were not informed about any potential previous treatment of the mice. Before analyzing the experimental outcome, investigators were blinded to the treatment of the mice. Therefore, the mouse numbers were allocated by lottery to random anonymous codes by one investigator. After replacing the original mouse numbers with the random codes on all relevant data files, the analysis was performed by a blinded second investigator. Only after receiving the numerical data (e.g. on infarct volumes, immune cells, etc.) were the anonymous codes changed back to the original mouse numbers.

### Statistical Analysis

Power analysis and sample size estimation were performed prior to the experiments using the G*Power V3.1.9.7 software (Heinrich Heine Universität Düsseldorf)[56]. Statistical analysis was performed using the Prism8 software (Graphpad). Data were tested for normal distribution by the Shapiro-Wilk or Kolmogorov-Smirnov test before applying parametric statistical tests. Two groups were compared using the unpaired Student’s t-test (parametric) or Mann-Whitney test (non-parametric), while multiple groups were compared by Kruskal-Wallis test following Dunn’s correction for multiple comparison due to small sample sizes that could not be tested for normal distribution. Data are shown as individual data points with mean and standard deviation (+/- SD) or standard error of the mean (+/- SEM). *= p < 0.05, **= p < 0.01, ***= p < 0.001, ****= p <0.0001. *Exclusion criteria*: A total of n = 7 mice (n = 5 for surgeon 1 and n = 2 for surgeon 2) did not survive 48 h post-tMCAO and therefore were excluded from further analysis (MRI & FACS).

## DATA AVAILABILITY

Sequence data reported in this publication have been submitted to the European Nucleotide Archive (ENA). The data is publicly available under accession PRJEB114334. Additional data will be shared by the corresponding author upon reasonable request.

## ACKNOWLEDGEMENTS

We thank Lennard Kuchenbecker-Pöhls, Karolin Gustmann, and Tobias Großhauser for their excellent technical assistance. We gratefully acknowledge the fruitful discussions with members of SFB1328 and the ENDomics & ERSI laboratories. We thank Carsten Bokemeyer, Director of the University Cancer Center Hamburg (UCC Hamburg), Hamburg, Germany, for his continuous support of the ENDomics Lab. We would like to thank the Cytometry and Cell Sorting Core Facility of the University Medical Center Hamburg-Eppendorf (UKE) for their excellent support with flow cytometry and data analysis. We also acknowledge the Forschungstierhaltung (FTH) of the UKE for their assistance in animal husbandry and care, which was essential for the completion of this work. Figure 1B (https://BioRender.com/1vqsc04) & Figure 3D (https://BioRender.com/yo2yyyg) were created in BioRender and are licensed under CC BY 4.0.

## FUNDING SOURCES

This study was supported by the German Research Foundation (DFG) in the context of SFB1328 (project number 335447717 to AW, BR, T Magnus & JK)

## AUTHOR CONTRIBUTIONS

BR, T Magnus & JK conceived and planned the study. AW, BR, T Magnus & JK guided the experiments. MS, EMC, JL, AV, KR & JH performed the laboratory experiments. MS performed the animal experiments. T Mummert & MK performed MRI measurements. CS & MA performed bioinformatic data analysis. BR, T Magnus & JK analyzed and interpreted data. MS & JK prepared the figures. MS, JK & T Magnus wrote the manuscript. JW, BR, T Magnus & JK revised the manuscript. All authors discussed the results and approved the final manuscript.

## ETHICAL APPROVAL

The animal experiment was conducted in accordance with the ARRIVE guidelines, the German Animal Welfare Act (TierSchG) and the European Union’s legislation on the protection of animals used for scientific purpose (EU directive 2010/63/EU). The ethical application and the study protocol were reviewed and approved by the responsible local authority (Behörde für Justiz und Verbraucherschutz der Freien und Hansestadt Hamburg - Lebensmittelsicherheit und Veterinärwesen) and by the animal welfare officers of the University Medical Center Hamburg-Eppendorf. Human primary cells (hBMEC) were obtained from ScienCell. All ScienCell Research Laboratories human cells are sourced within the USA under protocols that have obtained Institutional Review Board (IRB) approval, governed by Title 45 of the Code of Federal Regulations (CFR) Part 46. IRBs are federally empowered by the US Food and Drug Administration (FDA) and the Department of Health and Human Services who authorize and task them to review, modify, approve or disapprove proposed research involving human subjects under the regulations of 45 CFR Part 46. Fundamental to obtaining IRB approval is that the important ethical tenets outlined in the Belmont Report (Ethical Principles and Guidelines for the Protection of Human Subjects of Research, Report of the National Commission for the Protection of Human Subjects of Biomedical and Behavioral Research) are adhered to.

## CONFLICT OF INTEREST STATEMENT

JK is listed as an inventor on a patent on AAV-BR1 held by Boehringer Ingelheim GmbH.

## Notes

### Competing Interest Statement

Jakob Koerbelin is listed as an inventor on a patent on AAV-BR1 held by Boehringer Ingelheim GmbH.

