## Supplementary Figures for "CD73-derived adenosine at the blood-brain barrier confers protection in a mouse model of ischemic stroke"

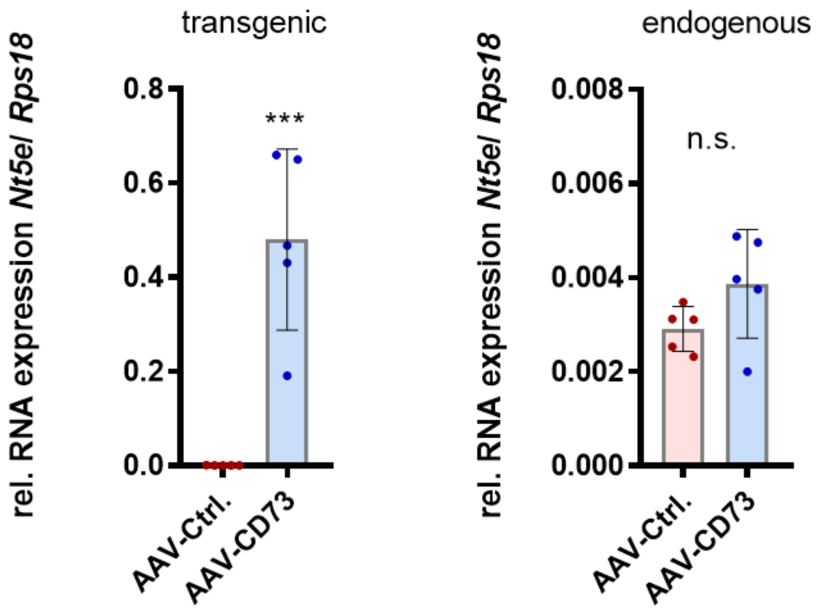


**Supplementary Figure 1. AAV-mediated CD73 expression in murine bEnd.3 cells.** Expression of transgenic (left) and endogenous (right) *Nt5e* (CD73*)* mRNA in the murine brain endothelial cell line bEnd.3 relative to the RPS18 housekeeping gene. Statistics: Two-tailed t-test; *** p = 0.0005; ns = not significant; n = 5 independent experimen

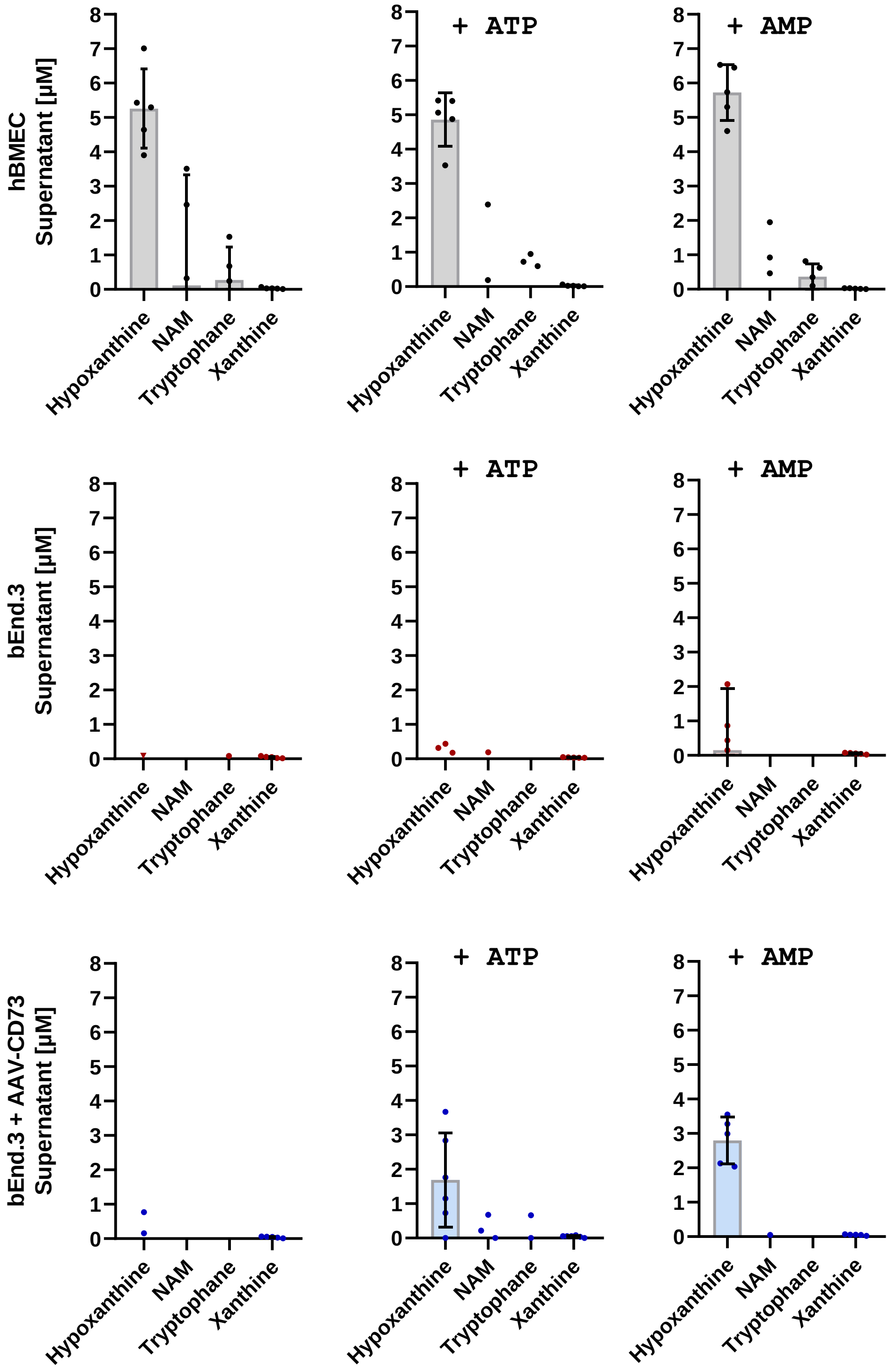


**Supplementary Figure 2**. The concentration of extracellular purine metabolites in the supernatant of primary human brain microvascular endothelial cells (hBMEC; upper panel), murine brain endothelial cells (bEnd.3; intermediate panel) and murine brain endothelial cells (bEnd.3) treated with AAV-BR1-CAG-CD73 vector (100,000 genomic particles/cell; lower panel) was determined by LC-MS/MS. Probes were collected 30 minutes after challenging the cells either with 100 µM ATP (middle) or 100 µM AMP (right) and compared to unchallenged cells (left). A total of n = 5 independent experiments were performed per cell type and treatment.

**CD73-derived adenosine at the blood-brain barrier is highly protective in a mouse model of ischemic stroke**

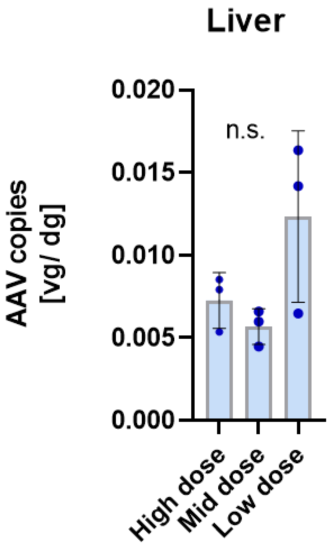


**Supplementary Figure 3. AAV vector copies in the liver.** Mice either received 5 x $10^{11}$ (low dose), 1 x $10^{12}$ (mid dose), or 2.5 x $10^{12}$ (high dose) genomics particles (gp)/ animal via the tail vein**.** The number of AAV vector copies (vg) per diploid mouse genome in the livers of n = 3 mice/ group was determined by qPCR 14 days after vector administration. Vector copies are indicated as viral genomes (vg) per diploid mouse genome (dg). Statistics: Kruskal-Wallis test; ns= not significant

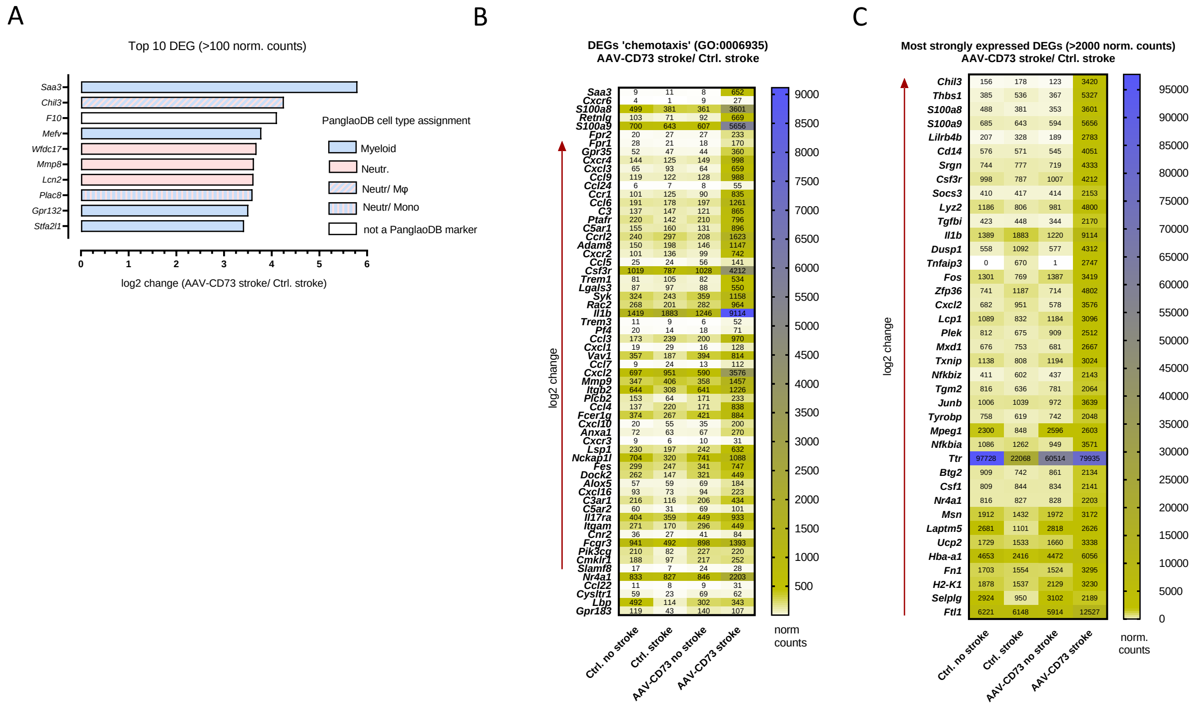


**Supplementary Figure 4. Differentially expressed genes in leukocytes of mice expressing *Nt5e* at the BBB. A**) According to PanglaoDB (https://panglaodb.se/), the top 10 most strongly regulated DEGs in CD45$^{+}$ brain cells were either mainly attributed to myeloid cells (blue bars), neutrophils (pink bars), or equally expressed by monocytes/macrophages and neutrophils (hatched blue/pink bars). **B**) Heatmap including all genes belonging to the GO term "chemotaxis" that were upregulated in CD45$^{+}$ cells from murine brains. **C**) Heatmap including the 40 most strongly expressed DEGs in CD45$^{+}$ cells from murine brains.
